# *CAMOR* and specific oncogene-driven lncRNAs mediate carcinogenic functions downstream of *MYC*, mutant *KRAS*, and mutant *TP53*

**DOI:** 10.64898/2026.09.04.749381

**Authors:** Maria Grześ, Akanksha Jaiswar, Wojciech Kaźmierczak, Tomasz Olesiński, Magdalena Nowak-Niezgoda, Dawid Walerych

**Affiliations:** Mossakowski Medical Research Institute PAS, Warsaw, Poland; Maria Sklodowska-Curie National Research Institute of Oncology, Warsaw, Poland; National Medical Institute of the Ministry of the Interior and Administration, Warsaw, Poland

**Keywords:** long non-coding RNA, oncogene regulation, CAMOR, RAKRAR, LINC00997, AC104447.1

## Abstract

**Purpose:** Long non-coding RNAs (lncRNAs) are important regulators of tumor biology, but their dependence on major oncogenic drivers and cancer specificity remain poorly characterized. We aimed to identify lncRNAs regulated by *MYC*, mutant *KRAS*, and mutant *TP53* across colorectal, lung, and pancreatic cancer and to determine their functional relevance and molecular mechanism.

**Methods:** We performed an integrative analysis of lncRNA expression profiles driven by *MYC*, mutant *KRAS*, and mutant *TP53* across colorectal, lung, and pancreatic cancer cell lines. Candidate lncRNAs were functionally characterized using assays of cancer cell viability, migration, and clonogenicity. Expression in patient-derived tumor and normal tissues and downstream molecular pathways were investigated.

**Results:** *MYC* was associated with the most extensive lncRNA regulation, with expression patterns largely independent of tissue origin. *MYC*-dependent *LINC00997* and the previously uncharacterized *AC104447.1,* as well as specifically KRAS-dependent *RAKRAR* (RASSF3 antisense KRAS regulated lncRNA) promoted cancer cell viability, migration, and clonogenicity. *LINC00997* and *AC104447.1* expression was elevated in specific tumor types compared to normal tissues. We identified *CAMOR* (CARNMT1 Antisense Multiple-Oncoprotein Regulated lncRNA), a pan-cancer lncRNA upregulated by all three oncogenic drivers. CAMOR promoted viability and migration specifically in cancer cells across analyzed cell lines and tumor types. Mechanistically, *CAMOR* and neighboring *CARNMT1* form a negative feedback regulatory loop affecting *CAMOR* oncogenic activity. Distinct downstream gene expression signatures suggested that the studied lncRNAs act through different regulatory processes.

**Conclusion:** Our findings expand the repertoire of oncogenic lncRNAs and demonstrate that integrating multiple oncogenic contexts enables identification of both driver-specific and broadly acting lncRNAs, providing a framework for identifying lncRNAs that may contribute to cancer development and represent potential diagnostic or therapeutic targets.

## INTRODUCTION

Major challenges in the management of cancer include late diagnosis, high recurrence rates, and the development of drug resistance (1, 2). This underscores an urgent need to identify novel biomarkers of cancer initiation and progression, markers for monitoring therapeutic efficacy, and new therapeutic targets. Lung, colorectal, and pancreatic cancers are among the six most lethal neoplasia types (1, 2), and are frequently driven by overexpression of *MYC*, and by oncogenic mutations in *KRAS* and *TP53* (3–6). Importantly, options to therapeutically target this trio of oncogenes are limited to several experimental and resistance-prone protocols (7–12). As a possible alternative we demonstrated in our recent work that cancers driven by c-MYC, mutant KRAS, or mutant p53 can be effectively targeted by inhibiting key downstream effectors (13). Finding these therapeutic opportunities was possible thanks to an analysis of mRNA and protein programs, while the transcriptomics data generated in a panel of cell lines included additional information on thousands of long non-coding RNAs driven by the trio of the studied oncogenes. In this study we present the results of their large-scale analysis and finding lncRNAs with functional oncogenic properties.

Long non-coding RNAs (lncRNAs) defined as non-coding transcripts longer than 200 nucleotides (14), emerged as critical components in tumor biology (15). LncRNAs have pivotal roles in cellular processes (including cell cycle, metabolism, apoptosis, and differentiation), gene expression regulation (on epigenetic, transcriptional, and translational level), and their aberrant expression is associated with tumorigenesis, tumor progression, and metastasis (16, 17). One lncRNA can be involved in the onset and progression of various neoplasias (18). Experimental data have shown that specific lncRNAs play an important role in the progression of pancreatic (19), lung (20), and colorectal (21) cancers, and represent possible therapeutic targets. However, among thousands of identified human lncRNAs, only a small fraction has been thoroughly functionally characterized, highlighting the need for further investigation (14). Unlike many mRNAs, the majority of lncRNAs exhibit tissue-specific expression patterns, while ubiquitous ones were also described (22, 23). One prominent example is metastasis-associated lung adenocarcinoma transcript 1 (MALAT1), a ubiquitously expressed lncRNA extensively studied for its roles in multiple cancers progression, therapy resistance, and potential as a therapeutic target (18, 24). Interestingly, MALAT1 role in normal cells has not been determined.

Accumulating evidence suggests that oncoproteins such as c-MYC, mutant KRAS, and mutant p53 can act as key regulators of lncRNA expression. For example MILIP, a c-Myc-Inducible lncRNA, functions as an inactivator of the tumor suppressor wild-type *TP53* (25). *HIF1A-As2*, a mutant KRAS-induced lncRNA, forms a double-positive feedback loop with c-Myc to promote proliferation and metastasis in NSCLC (non-small cell lung cancer) (26). Mutant p53 transcriptionally regulates MIR205HG, which contributes to proliferation, migration, and clonogenicity in head and neck squamous cell carcinoma cell lines (27). This research shows that lncRNAs are often found to act specifically in the context of cancer types and driver oncogenes, and ubiquitous regulators such as MALAT1 are in minority among the neoplasia-relevant lncRNAs which have been characterized (28).

In attempt to find both specific and universal oncogenic lncRNAs in this study we defined and overlapped c-Myc, mutant KRAS, and mutant p53-driven lncRNA expression profiles in a panel of colorectal and lung cancer cell lines. Through this approach, we identified and confirmed pro-oncogenic functional profile of two lncRNAs specifically dependent on c-Myc (*LINC00997* which was previously reported and *AC104447.1*, which is a novel one), one lncRNA, *RAKRAR* (RASSF3 antisense KRAS regulated lncRNA), selectively regulated by KRAS, and one lncRNA commonly activated by the studies oncogenes - *CAMOR* (CARNMT1 Antisense Multiple-Oncoprotein Regulated lncRNA) involved in a previously unrecognized interaction with CARNMT1 protein. Our findings expand current knowledge of oncogenic lncRNAs regulated by the major cancer drivers.

## MATERIAL AND METHODS

### Cell lines

A-549 (RRID:CVCL_0023), NCI-H23 (RRID:CVCL_1547), NCI-H1299 (RRID:CVCL_0060), DLD-1 (RRID:CVCL_0248), RKO (RRID:CVCL_0504), HT29 (RRID:CVCL_A8EZ), MIA PaCa-2 (RRID:CVCL_0428), Capan-2 (RRID:CVCL_0026), PANC-1 (RRID:CVCL_048), and BxPC-3 (RRID:CVCL_0186) cell lines were acquired from the American Type Culture Collection (ATCC, Manassas, VA, USA). LoVo (RRID:CVCL_0399) cell line was purchased from the European Collection of Authenticated Cell Cultures (ECACC, Salisbury, UK) repository. VMRC-LCD (RRID:CVCL_1787) cell line was acquired from Japanese Collection of Research Bioresources Cell Bank (JCRB, Ibaraki Osaka, Japan). The cells were routinely tested for the *Mycoplasma* sp. Presence by RT-qPCR. Only low passage numbers (below 15 post acquisition) of cell lines were used for the experiments.

Cell lines were cultured in an appropriate medium – RPMI (Gibco, Life Technologies, Rockville, MD, USA) for NCI-H23, NCI-H1299, DLD-1, Capan-2, and BxPC-3 and DMEM medium (Gibco) for A-549, RKO, HT29, MIA PaCa-2, PANC-1, LoVo, and VMRC-LCD, supplemented with 10% fetal bovine serum (FBS; Gibco) and 1% Pen Strep Antibiotics (Gibco). Cell culture was performed in a humidified atmosphere with 5% CO_2_ at 37°C. Human primary fibroblasts (F02 and F03) were obtained from skin biopsies of healthy subjects based on the Central Clinical Hospital of Ministry of Interior and Administration in Warsaw bioethics committee approval (Nos. 108/2017 and 203/2020) (29). Human h-TERT (telomerase reverse transcriptase) immortalized K15 fibroblasts were a kind gift from Prof. Harm Kampinga (University of Groningen, Netherlands) (30). Fibroblasts were cultured in DMEM medium (Gibco) supplemented with 10% FBS (Gibco) and 1% Pen Strep Antibiotics (Gibco).

### Bioinformatics analysis and RNA-seq data availability

Raw RNA-seq data generated in our previous research (13), deposited in the GEO database under accession number GSE239817, were analyzed. Raw reads were filtered and trimmed, mapped to the human reference genome, and transcript abundance was quantified according to the protocol described previously (31).

### Identification of lncRNAs

The pipeline developed in this study to identify lncRNAs was adapted, with minor modifications, from previously published methods (32). Transcript assembly was performed using Stringtie software with BAM files from each sample, followed by the merge function to generate a reference transcriptome (33). The assembled transcriptome was compared with the human reference genome (hg19) using the gffcomapre (34). Long noncoding RNA annotations were downloaded from Gencode (https://www.gencodegenes.org/human/) (35). Read counts for each gene were calculated using the FeaturesCount R package (36).

Transcripts which did not meet the following criteria were excluded: (i) length <200 bp; (ii) overlap of the database annotation with the exonic region; (iii) fragments per kilobase of transcripts per million mapped reads (FPKM) with values ≤ 0.5 and (iv) failure to pass the protein-coding score test by the Coding Potential Calculator (CPC), and Coding-potential assessment tool (CPAT2) (37, 38). Hierarchical clustering analysis was performed using Pearson correlation.

### Principal Component Analysis

Raw count data were normalized using size factors estimated with estimateSizeFactors. The geometric mean of counts and the median of ratios within each sample were calculated. Normalized expression data were used to generate principal component analysis (PCA) plots. The tidyverse package and the prcomp function in R were used for PCA visualization.

### Quantification of gene expression and identification DE lncRNAs

Mapped reads (BAM files) were processed using featureCounts to quantify transcript abundance and assess mapping quality. The resulting count matrix was analyzed using DESeq2 to identify differentially expressed lncRNAs (DElncRNAs) between normal and control conditions. Gene expression values were normalized using the Relative Log Expression (RLE) method implemented in DESeq2. Adjusted P-values were calculated using the Benjamini–Hochberg method. Transcripts with adjusted P < 0.05 and log2 fold change ≥ 2.0 were considered significantly differentially expressed lncRNAs.

### Patient’s samples

In this study, tumor tissue, adjacent normal tissue, and peripheral blood samples from 10 patients with colon cancer and 11 patients with pancreatic cancer undergoing surgical treatment were used. Histopathology characterization of the samples and confirmation of their cancer status was shown in our previous papers (13, 31). The study was approved by the Bioethics Committees of the National Medical Institute of the Ministry of the Interior and Administration in Warsaw, Poland (Approval No. 109/2016, with subsequent amendments) and the National Institute of Oncology in Warsaw, Poland (Approval No. 55/2023). Written informed consent for the research use of collected tissues was obtained from all patients. Immediately after surgical resection, tissue samples were snap-frozen in liquid nitrogen and stored until further analysis. Blood samples were stored at −80°C for subsequent use.

### Gene expression silencing

Cell transfections were performed using Lipofectamine™ RNAiMAX (Invitrogen, ThermoFisher Scientific, Waltham, MA, USA) according to the manufacturer’s instructions. siRNAs targeting the analyzed lncRNAs (provided by Sigma-Aldrich, St. Louis, MO, USA) and a scrambled siRNA used as a negative control are listed in Supplementary Table 3. Specificity of each siRNA was checked by qPCR. Cells were transfected twice, with the second transfection performed 24 hours after the first. Subsequent experiments were conducted 48 hours after the second transfection. Each siRNA was used at a final concentration of 20 nM.

### Total RNA extraction from cell lines and patient samples

Total RNA was extracted from cultured cells and patient tissue samples utilizing RNA Extracol (EURx, Gdansk, Poland), following standard phenol-chloroform RNA isolation protocol., Patient tissue specimens were homogenized in RNA Extracol using RNA extraction beads (Diagenode, Liège, Belgium) during sonication using the Bioruptor® Plus system (Diagenode), followed by the standard phenol–chloroform RNA isolation procedure RNA isolation. Blood samples were first treated with RBC lysis buffer to remove erythrocytes. The resulting white blood cell pellet was subsequently processed with RNA Extracol following the standard protocol. RNA concentration and purity were assessed using a NanoDrop spectrophotometer (Thermo Fisher Scientific). Isolated RNA was treated with DNase I (A&A Biotechnology) to eliminate residual genomic DNA contamination.

### RT-qPCR

For cDNA synthesis, 500 ng of total RNA was reverse transcribed using the NG dART RT Kit (EURx) according to the manufacturer’s instructions. Quantitative PCR (qPCR) was performed using qPCR Sensitive RT HS-PCR Mix SYBR (A&A Biotechnology, gdynia, Poland) on a CFX Maestro™ Real-Time PCR System (Bio-Rad, Hercules, CA, USA). Relative gene expression levels were calculated using the 2^−ΔΔCt method, with GAPDH serving as the endogenous reference control. Primer sequences used for qPCR are listed in Supplementary Table 3.

### Droplet digital PCR (ddPCR)

Droplet digital PCR workflow was performed using our previous protocol (39), using a 20 μL volume of PCR mix containing 10 μL of 2X QX200 ddPCR EvaGreen Supermix (Bio-Rad), 6 μL of nuclease-free water and 4 μL of diluted cDNA. Each ddPCR assay mixture and 70 μL of droplet generation oil for EvaGreen (Bio-Rad) was loaded into a DG8 Cartridge (Bio-Rad), which was placed inside the QX200 Droplet Generator (Bio-Rad). Upon completion of droplet generation, the droplets were transferred to a 96-well PCR plate, which was placed in a thermal cycler. Thermal cycling conditions were: 95°C for 5 min, then 40 cycles of 95°C for 30 s and 60°C for 1 min and three final steps at 4°C for 5 min, 90°C for 5 min and a 4°C infinite hold with a ramping rate of 2°C/second in every step. Next, the sealed plate was transferred into the QX200 Droplet Reader (Bio-Rad). Obtained data were analyzed with QuantaSoft software v1.7.

### Subcellular fractionation

Cytoplasmic and nuclear RNA fractions were isolated using the Cytoplasmic and Nuclear RNA Purification Kit (Norgen Biotek, Thorold, ON, Canada), according to manufacturer’s manual. Briefly, cells cultured in 10-cm dishes were harvested and centrifugated, and the cytoplasmic fraction was carefully separated from nuclear pellet. RNA from both fractions was subsequently extracted following standard phenol-chloroform RNA isolation protocol. The subcellular localization of the analyzed lncRNAs was assessed by RT-qPCR method. GAPDH was used as a cytoplasmic reference transcript, while U6 served as a nuclear reference transcript.

### Cell viability assay and drug treatment

A total of 1 × 10^4^ cells were seeded in 90 µl of culture medium per well in 96-well plates. Forty-eight hours after the siRNA transfection/drug treatment, cells were washed with PBS, and ATPlite reagent (PerkinElmer, Waltham, MA, USA) was added according to the manufacturer’s instructions. Luminescence was measured using an Infinite plate reader (Tecan). Standard chemotherapy regimens were tested for each cancer type: FOLFOX and FOLFIRI for colorectal cancer; FOLFIRINOX or paclitaxel combined with gemcitabine for pancreatic cancer; and cisplatin in combination with vinorelbine, gemcitabine, or paclitaxel (manufacturers/providers) for lung cancer.

### Wound healing assay

Cell migration was evaluated using a wound healing (scratch) assay. Twenty-four hours after the second siRNA transfection, 3 × 10⁵ cells were seeded into 12-well plates and incubated overnight. Once cells reached 100% confluence, a linear scratch was created in the monolayer using a 200-µL pipette tip. Detached cells were removed by washing with PBS. Images of wound closure were captured at 0 h and after 24 h or 48 h using a light microscope. The wound area was quantified using ImageJ software with the Wound Healing Size Tool (updated version) plugin.

### Colony formation

To evaluate colony-forming activity, cells were transfected with scrambled siRNA or siRNA targeting the indicated lncRNA, and 500 cells were subsequently seeded into six-well plates in complete growth medium. After 2 weeks of incubation, allowing the colonies to develop, cells were washed with PBS, fixed with 4% paraformaldehyde, and stained with 0.1% crystal violet for 30 min. Colonies were then washed with PBS, photographed, and quantified using the ImageJ software plugin.

### Lentivirus production and transfection

Lentiviral plasmids for *CARNMT1-AS1* and *CARNMT1* overexpression were purchased from VectorBuilder (Chicago, IL, USA). HEK293T cells (ATCC) were grown in 10-cm dishes to approximately 70% confluence, were transfected using 90 µg of polyethyleneimine (PEI; Sigma Aldrich) with two second-generation packaging plasmids - 5 µg pMD2.G and 10 µg psPAX2 and 15 µg of target lentiviral vectors carrying: CARNMT-1 or CARNMT coding sequences. Details of lentivirus production and transfection are described elsewhere (13).

### Chromatin immunoprecipitation

Chromatin immunoprecipitation (ChIP) was performed using the SimpleChIP Plus Sonication Chromatin IP Kit (Cell Signaling Technology, Danvers, MA, USA), according to the manufacturer’s instructions. PCR primers for predicted c-Myc and mutant p53 binding regions in the CARNMT1-AS1 promoter were designed based on ChIP-Atlas (https://chip-atlas.org/) analysis and are listed in Supplementary Table 3. Promoter occupancy was calculated using the Fold Enrichment Method (2 − ΔΔCt method).

### Western blot

Western blot experiment was conducted according to protocol described elsewhere (13). This list of the used primary antibodies is included in the Supplementary Table 3.

### Identification of Differentially Expressed Genes (DEGs) and Functional Enrichment Analysis

To investigate the downstream functional implications of LINC00997, AC104447.1, RAKRAR, and CAMOR expression in the pancreatic, lund, and colorectal TCGA cohort, patient samples were stratified into high- and low-expression groups based on the expression levels of the analyzed lncRNA. Differential gene expression analysis between the two groups was performed using the TCGAbiolinks and DESeq2 packages within the R environment based on SummarizedExperiment objects. To focus on transcriptionally upregulated pathways associated with high lncRNA expression, differentially expressed genes (DEGs) were filtered to retain only upregulated transcripts with a statistical significance threshold of p < 0.01 and a log fold change > 0. Genes meeting these criteria across pancreatic, lund, and colorectal cancer types were intersected, and only the common genes were carried forward for downstream analysis. Subsequently, functional enrichment analysis (GSEA) was conducted. The top 10 enriched pathways, ranked by the highest number of associated genes, were visualized using a dot plot generated with the ggplot2 package, illustrating the enrichment score/rich factor alongside p-values.

### Statistical analysis and data visualization

GraphPad Prism 8.0 was used for statistical analysis and data visualization. Experimental data are presented as the mean ± standard deviation (SD) or standard error of the mean (SEM), as indicated. P values <0,05 were considered statistically significant (*p<0,05, **p<0,01, ***p<0,001). Each figure legend specifies the number of replicates and the statistical test applied. All schematics were created using BioRender.com.

## RESULTS

### Identification of oncogene–specific and commonly regulated lncRNAs

To identify long non-coding RNAs (lncRNAs) whose expression is controlled by the major driver oncogenes, we utilized datasets previously obtained and published by our laboratory in the context of mRNA and protein analysis (13). These datasets were obtained from CRISPR-Cas9-mediated downregulation of *MYC,* mutant *KRAS*, and mutant *TP53* expression in a panel of eight lung and colon cancer cell lines (Fig. 1A). The panel was designed to include for each cancer type one cell line driven by all three oncogenes and one cell line driven by each individual oncogene. As described in (13) and shown in Fig. 1A, the expression of the driver oncogenes in each cell line was downregulated by introducing sgRNAs into cells with stable Cas9 expression. After 48 hours, samples were collected in triplicates and RNA sequencing was performed. For the purpose of this study, a dedicated analysis was performed, focused on lncRNAs identification, dataset overlap, and functional validation of selected lncRNAs.

**Figure 1.**
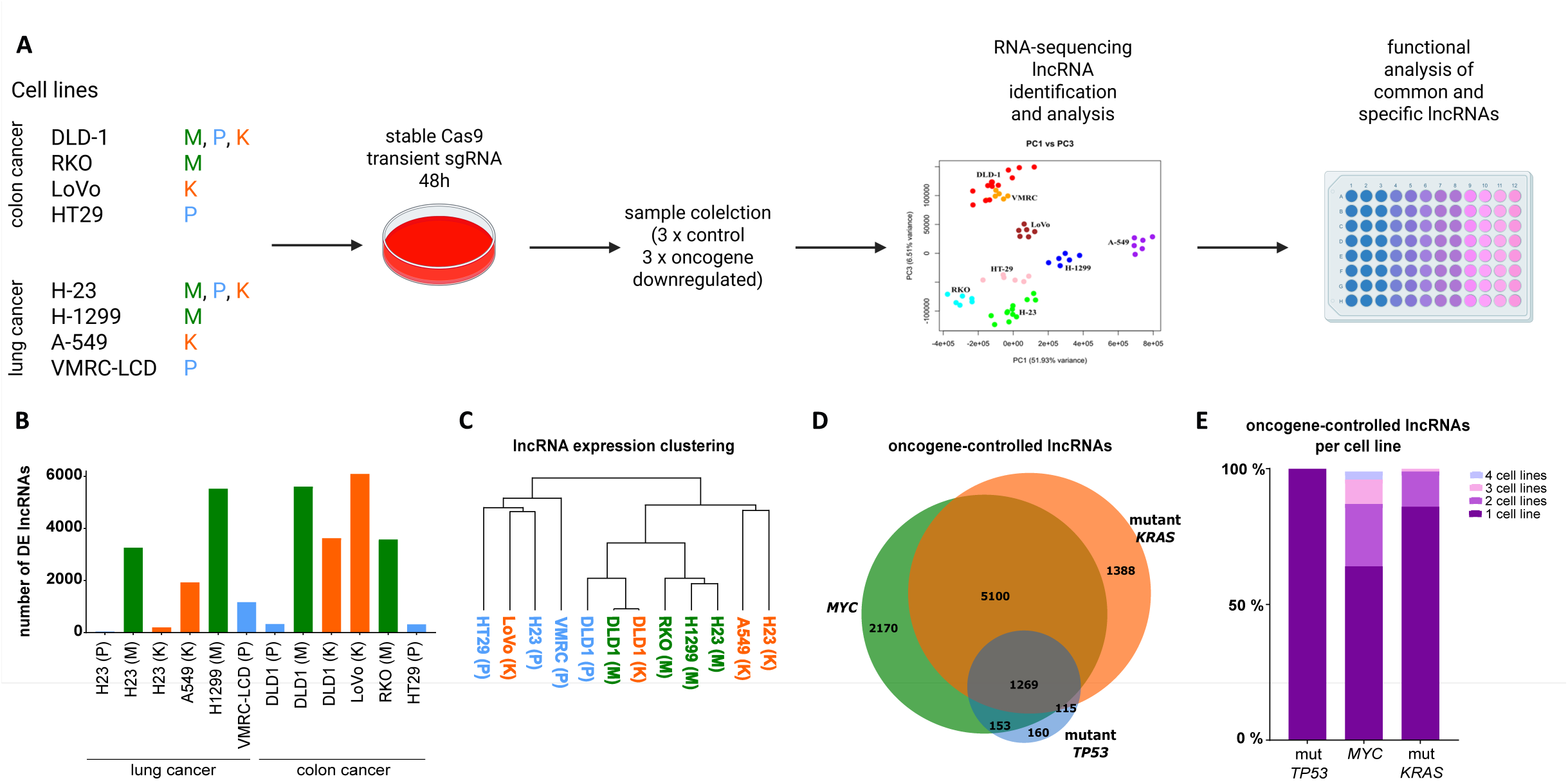
Experimental workflow and identification of differentially expressed lncRNAs. (A) Schematic overview of the workflow for CRISPR-mediated oncogene editing (M – *MYC*, P – mutant *TP53*, K – mutant *KRAS*) in the indicated cell lines, followed by lncRNAs bioinformatics and functional analyses. (B) Number of identified differentially expressed (DE) lncRNAs in each cell line, colors and letters in parentheses indicate the targeted oncogene. (C) Hierarchical clustering of the 6,845 lncRNAs common to all analyzed samples. Colors and letters in parentheses indicate the respective oncogene. (D) Venn diagram showing the overlap of DE lncRNAs regulated by the indicated oncogenes. (E) Bar plot showing the proportion of DE lncRNAs that are specifically regulated by a single oncogene across different numbers of cell lines.

Quality of RNA sequencing the data was assessed using PCA, (Suppl. Fig. 1A), identified lncRNAs were classified by their length (Suppl. Fig. 1B) and chromosomal distribution (Suppl. Fig. 1C). Differential expression analysis between control and samples with knocked-down oncogene revealed the highest number of differentially expressed lncRNAs (DElncRNAs) following *MYC* depletion, whereas mutant *TP53* downregulation resulted in the fewest changes in the lncRNA expression (Fig. 1B, Suppl. Fig. 1D–E). Hierarchical clustering of the DElncRNAs demonstrated that the samples with *MYC* downregulation cluster together, while samples with mutant *TP53* and mutant *KRAS* downregulation demonstrated less consistency in clustering (Fig. 1C).

We identified 1269 DElncRNAs commonly regulated by c-Myc, mutant KRAS, and mutant p53 (Fig. 1D, Suppl. Table 1). Additionally, we identified 2170 DElncRNAs specific to c-Myc, 1388 specific to mutant KRAS, and 160 specific to mutant p53. Analysis of the distribution of these DElncRNAs among cell lines revealed that all of the exclusively mutant p53-dependent lncRNAs were derived from single cell lines (one from H23, two from HT29, and 157 from VMRC), two mutant KRAS-specific lncRNAs were shared by three cell lines, and 74 c-Myc-specific DElncRNAs were detected across all four cell lines analyzed for c-Myc dependence (Fig. 1E). Furthermore, redundancy analysis, based on a methodology previously used by us for mRNAs (1), showed that the mutant p53 group contained the highest proportion of lncRNAs redundant to two other oncoproteins, while the c-Myc group consisted predominantly of non-redundant lncRNAs (Suppl. Fig. 1F-G). The majority of mutant p53 and KRAS redundant lncRNAs were controlled by c-Myc.

The above analyses suggested that c-Myc is the broadest and most consistent lncRNA expression regulator compared to other studied oncoproteins. Mutant p53 transcriptional programs had least specifically regulated lncRNAs and were most redundant. Hence we proceeded with detailed assessment of oncogenic expression and functions of c-Myc- and mutant KRAS-specific lncRNAs, followed by lncRNAs controlled by all studied oncoproteins.

### *LINC00997* and *AC104447.1* are oncogenic, specifically c-Myc-driven lncRNAs

Bioinformatics analysis revealed that c-Myc, compared to the other two analyzed oncogenes, significantly affects the expression levels of the highest number of lncRNAs. Moreover, a substantial proportion of these expression changes were specific to c-Myc, meaning that the expression of these lncRNAs remained unchanged following the downregulation of mutant *KRAS* or mutant *TP53* (Fig. 2A). In total, 26 lncRNAs were identified that met all three of the following criteria: upon CRISPR-Cas9-mediated *MYC* disruption their expression levels were downregulated (i) exclusively, (ii) consistently, and (iii) across all four cell lines tested upon *MYC* downregulation. For each of these lncRNAs, the average of expression level across the four cell lines was calculated, and the 10 lncRNAs with the lowest average were considered for further studies. Two lncRNAs—*AC104447.1* and *LINC00997* were selected for detailed analysis based on a positive validation by qPCR following *MYC* knockdown, using siRNA (Fig. 2A, Suppl. Table 2, Suppl. Fig. 2A-B). The expression levels of these two lncRNAs were found to be increased in K15, non-transformed, human immortalized fibroblasts (30) with lentiviral vector-mediated overexpression of *MYC* (Suppl. Fig. 2A-B). Analysis of a patient-derived data from The Cancer Genome Atlas (TCGA) showed that *LINC00997* is more highly expressed in lung and colorectal tumors compared to corresponding normal tissues (Fig. 2B). A similar pattern was observed for *AC104447.1* in colorectal tumors (Fig. 2C). In the case of pancreatic cancer, a statistically significant comparison could not be performed due to the limited number of normal tissue samples (n=4) available in the database. Thus, we interrogated in-house patient samples of pancreatic and colorectal cancers. For both lncRNAs studied, paired analysis comparing cancer and normal tissues revealed higher lncRNA expression in cancer samples (Fig. 2D-E). For this reason, we decided to include two pancreatic cancer cell lines, PANC1 and MIAPaCa-2, in further functional studies.

**Figure 2.**
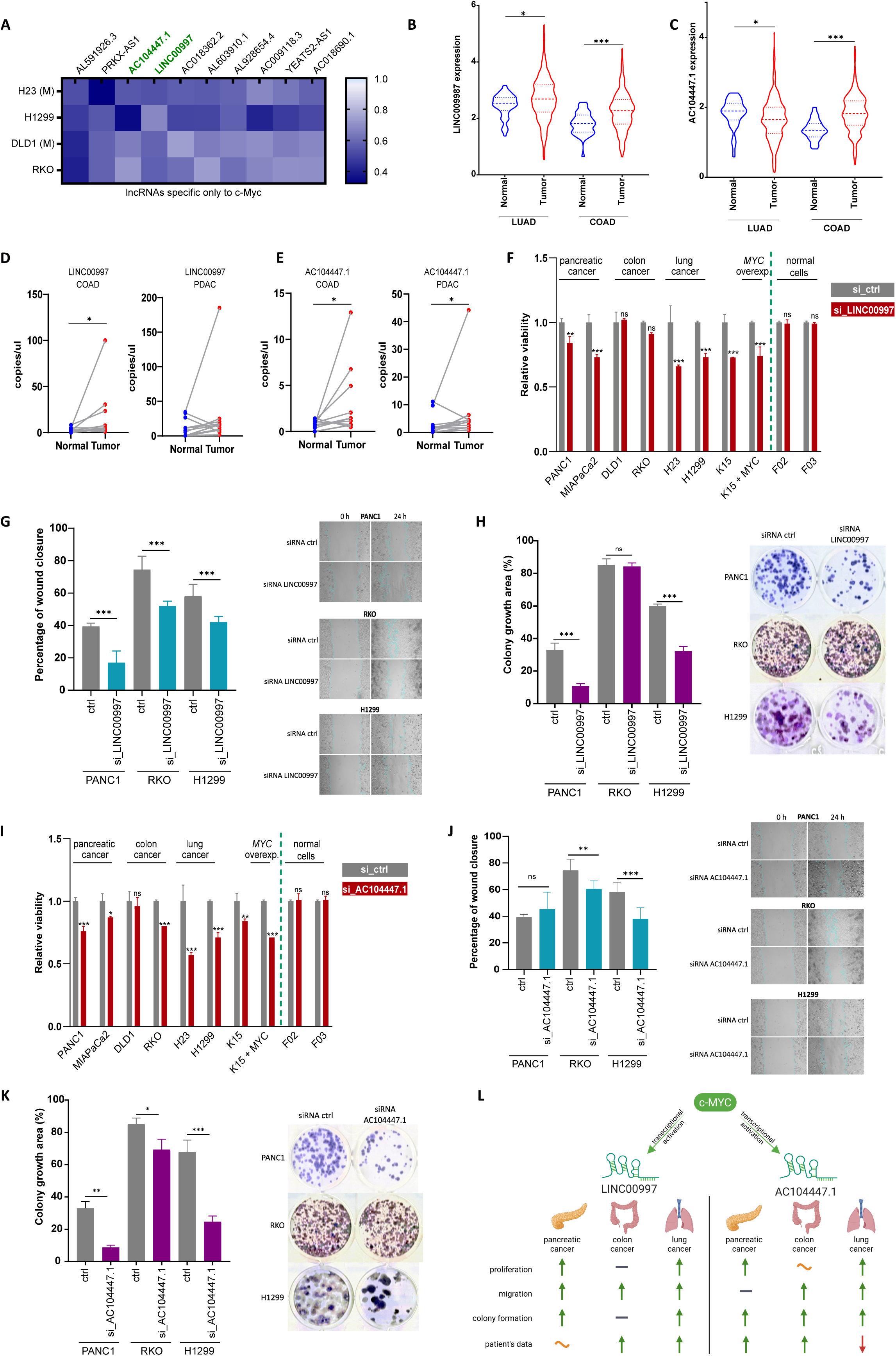
Identification of *LINC009997* and *AC104447.1*, *MYC*-specific lncRNAs, and their phenotypic analysis. (A) Heat map showing fold change (FC; range indicated on the color scale) of the top 10 *MYC* – specific lncRNAs exhibiting the strongest downregulation following *MYC* editing. TCGA data showing expression levels in colon and lung cancer versus normal tissues of (B) *LINC00997* and (C) *AC104447.1*. Data were analyzed with one-way ANOVA (uncorrected Fisher’s LSD). Expression of *LINC00997* (D) and *AC104447.1* (E) in paired colon (n=10) and pancreatic (n=11) cancer tissues compared to adjacent normal tissue, measured by ddPCR (Wilcoxon matched-pairs signed rank test was applied). (F) Effect of *LINC00997* silencing on the viability of pancreatic (PANC1, MIAPaCa2), colon (DLD1, RKO), and lung (H23, H1299) cancer cell lines, as well as normal fibroblasts (F02, F03) and fibroblasts overexpressing *MYC* (K15). Cell viability was measured using the ATPlite assay 48 h post-transfection. Data are presented as mean ± SD from *n* = 2 replicates per cell line and were analyzed using two-way ANOVA (uncorrected Fisher’s LSD) versus the siRNA negative control. (G) Cell scratch (wound healing) assay detecting the impact of *LINC00997* silencing on PANC1, RKO, and H1299 cells migration. Data were analyzed with one-way ANOVA (uncorrected Fisher’s LSD), n= 3 to 5 replicares. (H) Cell colony formation was conducted to evaluate the proliferative ability following *LINC00997* silencing. Data were analyzed with one-way ANOVA (uncorrected Fisher’s LSD) with n = 3 for each group. (I) Viability of pancreatic (PANC1, MIAPaCa2), colon (DLD1, RKO), and lung (H23, H1299) cancer cell lines, as well as normal fibroblasts (F02, F03) and fibroblasts overexpressing *MCY* (K15) following *AC104447.1* silencing. Data were presented and analyzed as in (F). Assessment of (J) migratory and (K) proliferative phenotypes in PANC1, RKO, and H1299 cell lines following *AC104447.1* silencing. (L) Schematic overview of the oncogenic roles of *LINC00997* and *AC104447.1*. B, C, D, F, G, H: \**p* < 0.05, \*\**p* < 0.01, \*\*\**p* < 0.001.

Both selected lncRNAs were found to exhibit predominantly nuclear localization (Suppl. Fig. 2C-D), which is considered a marker of potential transcription regulation activity (40). Functional analysis of *LINC00997* showed that its knockdown using siRNA reduced cell viability in pancreatic and lung cancer cell lines, while having no significant effect on the viability of colorectal cancer cells or normal fibroblasts (Fig. 2F). Further functional assays were narrowed to three cell lines representing each cancer type - H1299 (lung cancer), RKO (colon cancer), and PANC1 (pancreatic cancer). Knockdown of *LINC00997* inhibited migration of all these cell lines (Fig. 2G) as well as colony formation in H1299 and PANC1 (Fig. 2H). Additionally, correlation analysis of *LINC00997* expression levels in blood and pancreatic tumors from patients revealed a strong positive correlation (R = 0.71) in contrast with low correlation with normal margin tissue (R=-0.02) (Suppl. Fig. 2E).

The second potentially oncogenic lncRNA regulated by c-Myc was *AC104447.1*. Similarly, knockdown of this lncRNA led to decreased cell viability in pancreatic and lung cancer cell lines, as well as in one colorectal cancer line (RKO), without affecting the viability of normal fibroblasts (Fig. 2I). Knockdown of *AC104447.1* reduced the migratory capacity of H1299 and RKO cells (Fig. 2J) and impaired colony formation in H1299, DLD1, and PANC1 cell lines (Fig. 2K).

To identify the potential signaling pathways and biological processes through which *LINC00997* and *AC104447.1* may exert their oncogenic functions, we performed functional enrichment analysis on the set of upregulated DEGs identified across the pancreatic, lung, and colorectal cancers TCGA datasets. For *LINC00997* GSEA revealed significant enrichment of pathways associated with mitochondrial function and cellular energy metabolism, as well as gene sets involved in membrane and transmembrane transport (Suppl. Fig. 2F). In contrast, for *AC104447.1*, we observed enrichment in EGF/EGFR signaling pathway, focal adhesion, protein serine/threonine kinase activity, and cell–cell signaling (Suppl. Fig. 2G).

To experimentally validate the GSEA results, we analyzed the expression of NDUFV2 and RAC1 following *LINC00997* silencing (Suppl. Fig. 2H), and MAPK1 and ARHGAP5 after *AC104447.1* silencing (Suppl. Fig. 2I). *LINC00997* silencing, contrary to our initial expectation, significantly increased NDUFV2 expression in RKO and H1299 cell lines, which may reflect a compensatory or adaptive response to lncRNA depletion, potentially involving the activation of alternative metabolic or signaling pathways. In turn, ARHGAP5 transcript levels were significantly reduced in both PANC1 and H1299 cells following *AC104447.1* silencing, suggesting that *AC104447.1* may positively regulate ARHGAP5 expression.

In conclusion, both studied lncRNAs displayed promotion of cancer-associated phenotypes in selected cancer cell lines (Fig. 2L), which together with the patient-derived data suggested their pro-neoplastic activity and a diagnostic potential.

### Mutant *KRAS*-dependent lncRNA, *RAKRAR* impacts cancer cells viability, migration, and colony formation

Among the lncRNAs whose expression was altered following specifically mutant *KRAS* knockdown, only two candidates— one that we now call *RAKRAR* (RASSF3 antisense KRAS regulated lncRNA, formerly known as *AC078962.3*) and *SPRY4-AS1*—fulfilled the criteria of being consistently downregulated exclusively in response to KRAS silencing and being detected in multiple cell lines (specifically, in three cell lines) (Fig. 3A, Suppl. Table 2). Other 6 lncRNAs shown in Fig. 3A were mutant KRAS-regulated in four cell lines, but their expression changed also upon depletion of other studied oncogenes. Although qPCR validation confirmed the downregulation of *SPRY4-AS1* upon mutant *KRAS* knockdown, we failed to design an siRNA efficiently downregulating this transcript in cell lines (not shown). Therefore, further analysis focused on *RAKRAR* (Suppl. Fig. 3A), another nuclear-localized lncRNA (Suppl. Fig. 3B). Expression analysis of TCGA patient-derived datasets was not feasible due to a limited number of normal samples with detectable *RAKRAR* expression (n=8 in lung and colon cancers). Analysis of *RAKRAR* expression levels in normal and tumor tissues using the TCGA dataset was not feasible, as expression data for this lncRNA in normal lung and colon tissues were available for only eight normal samples. Furthermore, analysis of in-house patient cohorts did not reveal a statistically significant difference between paired normal and tumor tissues in colon and pancreatic samples (Fig. 2B). Functional assays demonstrated that silencing of *RAKRAR* reduced cell viability in lung, pancreatic, and colorectal cancer cell lines. Additionally, viability was also affected in one of the tested normal fibroblasts (Fig. 3C). Further functional validation revealed that *RAKRAR* knockdown impaired the migratory capacity of the pancreatic cancer cell line PANC1 (Fig. 3D) and negatively affected colony formation in A549 (lung cancer), DLD1 (colorectal cancer), and PANC1 cell lines (Fig. 3E).

**Figure 3.**
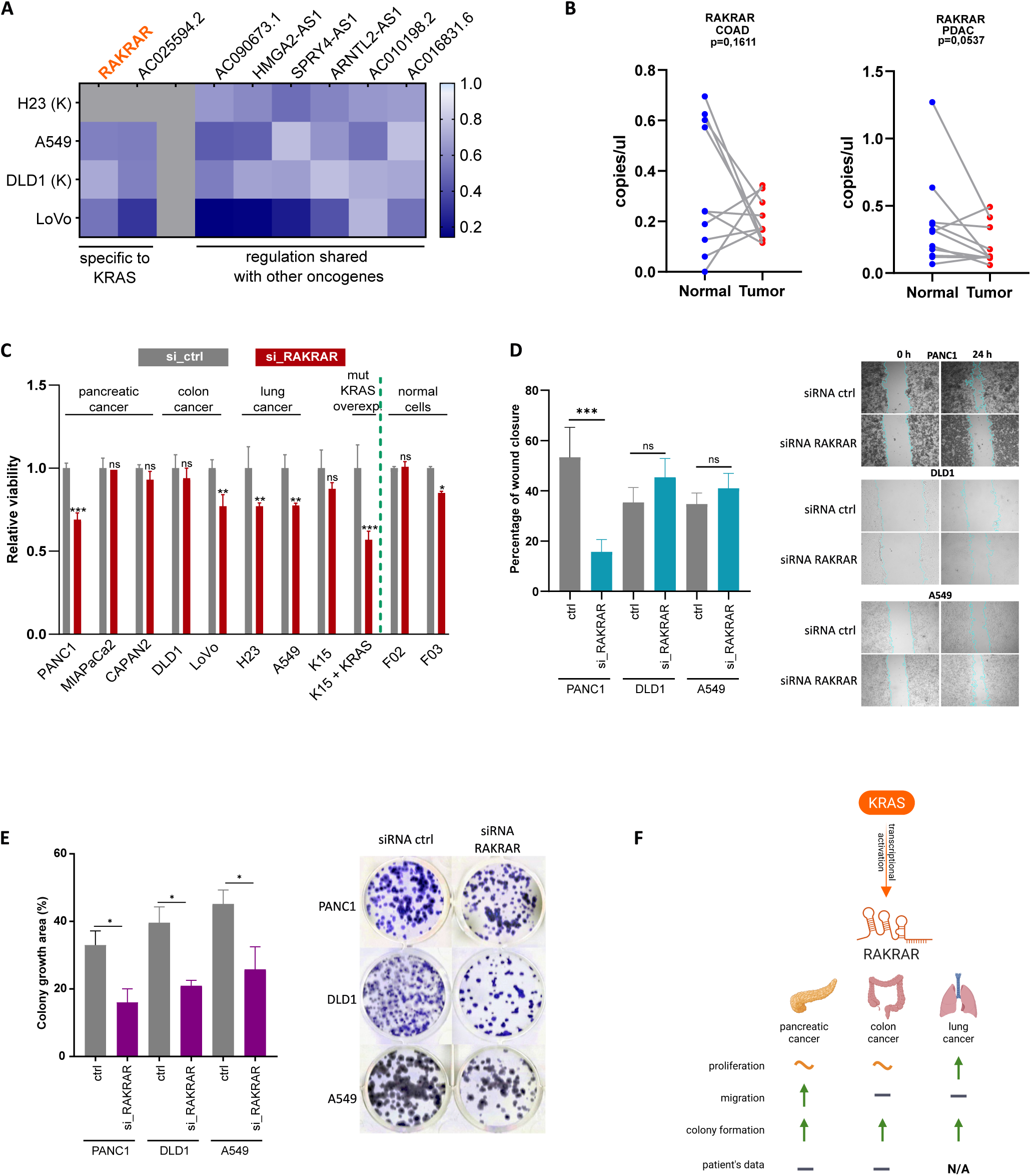
*RAKRAR* acts as a mutant *KRAS*-dependent pro-oncogenic lncRNA. (A) Heat map showing fold change (range indicated on the color scale) of two mutant KRAS-specific and six non-specific lncRNAs exhibiting the strongest downregulation following *KRAS* editing. (B) ddPCR analysis of *RAKRAR* levels in colon (n=10) and pancreatic (n=11) cancer tissues and their paired adjacent non-cancerous tissues. (C) Effect of *RAKRAR* silencing on the viability of pancreatic (PANC1, MIAPaCa2, CAPAN2), colon (DLD1, LoVo), and lung (H23, A549) cancer cell lines, as well as normal fibroblasts (F02, F03), and fibroblasts overexpressing *KRAS* (K15). ATPlite assay was applied to measure cell viability 48 h post-transfection and two-way ANOVA (uncorrected Fisher’s LSD) was used to analyze data versus the siRNA negative control. (D) Scratch (wound healing) assay assessing the impact of *RAKRAR* silencing on PANC1, DLD1, and A549 cells. Data were analyzed using one-way ANOVA (uncorrected Fisher’s LSD), *n* = 3–5 replicates. (E) Colony formation assay of PANC1, DLD1, and A549 cells with silenced *RAKRAR*. Data were analyzed using one-way ANOVA (uncorrected Fisher’s LSD), *n* = 3 per group. (F) Schematic overview of the oncogenic functions of *RAKRAR*

For *RAKRAR*, GSEA revealed significant enrichment of gene sets associated with cell adhesion, regulation of cell adhesion, and transmembrane signaling receptor activity (Suppl. Fig. 3C). To validate these findings, the expression of MAD1L1 and CXCL13 was analyzed following *RAKRAR* silencing. Interestingly, *RAKRAR* depletion significantly reduced CXCL13 transcript levels in PANC1 and DLD1 cells, with a similar trend observed in A549 cells (Suppl. Fig. 3D). Obtained results suggested the pro-oncogenic role of *RAKRAR* in selected cancer cell lines, while have not confirmed its upregulation in cancer vs. normal tissues (Fig. 3F).

### *CAMOR* is an oncogenic lncRNA driven redundantly by mutant *KRAS*, mutant *TP53*, and *MYC*

Among the lncRNAs whose expression was altered following the downregulation of all three oncogenes (*MYC*, mutant *KRAS*, and mutant *TP53*), five lncRNAs met the criteria of consistent downregulation and detection in at least nine out of ten analyzed cancer cell lines (Fig. 4A). Out of these, an lncRNA that we now designate as *CAMOR* (CARNMT1 Antisense Multiple-Oncoprotein Regulated, also known as *CARNMT1-AS1*), emerged as the most promising candidate for functional characterization based on its unknown role in cancer and a positive validation by qPCR following *MYC*, mutant *KRAS*, and mutant *TP53* knockdown using siRNA (Suppl. Table 2, Suppl. Fig. 4A). Analysis of TCGA patient-derived data revealed higher expression of *CARNMT1-AS1* in colon and lung tumors compared to corresponding normal tissues (Fig. 4B). Similarly, in-house patient samples of pancreatic (n=8) and colorectal (n=9) cancers analysis comparing cancer and normal tissues revealed higher lncRNA expression in cancer samples (Fig. 4C). Moreover, correlation analysis of *CAMOR* expression levels in blood and pancreatic tumors from patients revealed a significant positive correlation (R = 0.6) in contrast with insignificant correlation with normal margin tissue (Suppl. Fig. 4B).

**Figure 4.**
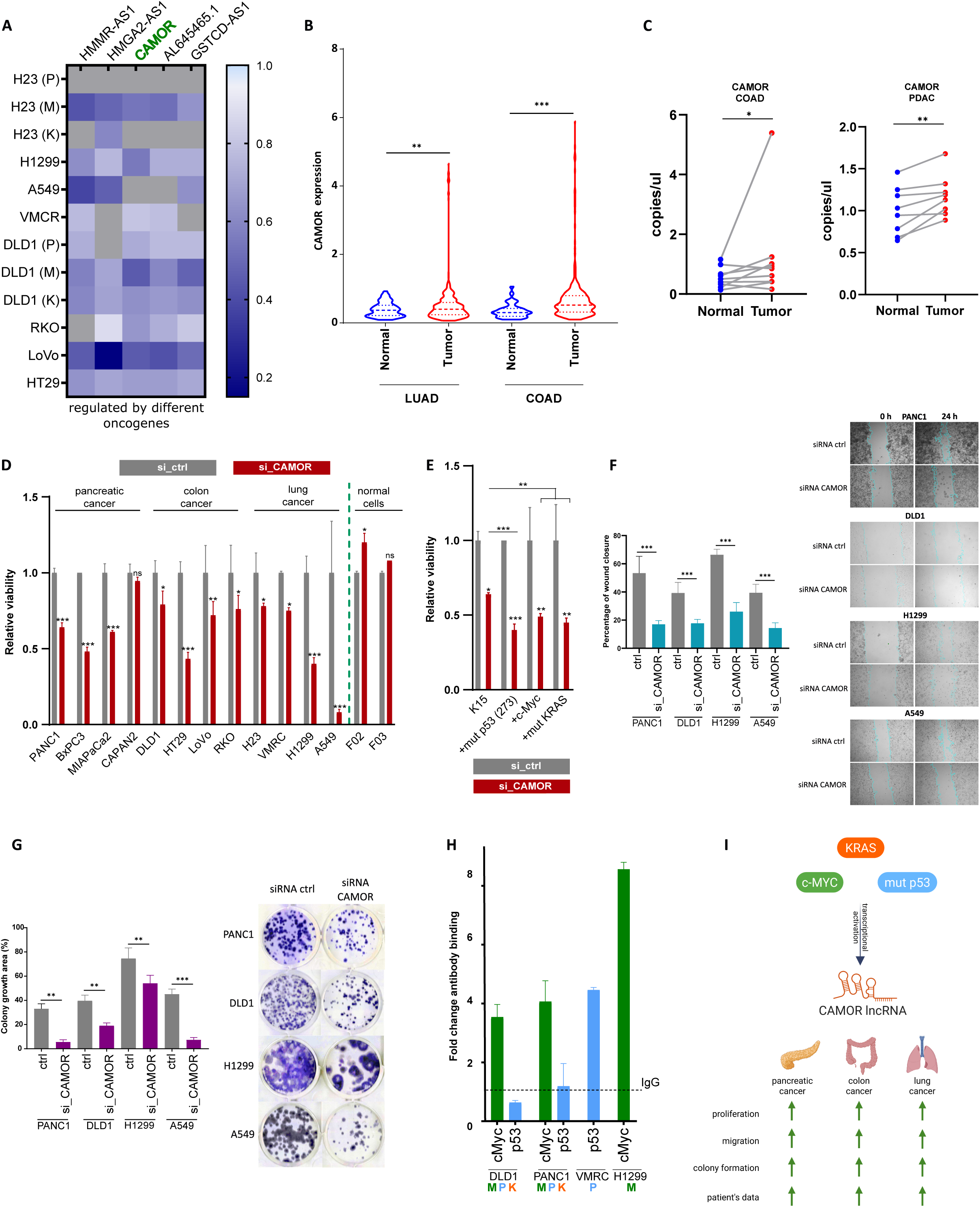
Pro-oncogenic effects of *CAMOR*, a pan-cancer lncRNA regulated by all three oncogenic drivers. (A) Heatmap illustrating fold changes (scale indicated) of the five most consistently across at least 9 cell lines lncRNAs that show the most pronounced downregulation upon *MYC*, mutant *KRAS*, and mutant *TP53* editing. (B) TCGA-derived expression levels of CAMOR in colon and lung cancer tissues compared with corresponding normal tissues. (C) Expression of *CAMOR* in paired colon (n=9) and pancreatic (n=8) cancer tissues compared to adjacent normal tissue, measured by ddPCR (Wilcoxon matched-pairs signed rank test was applied). (D) Impact of *CAMOR* knockdown on cell viability in pancreatic (PANC1, BxPC3, MIAPaCa2, CAPAN2), colon (DLD1, HT29, LoVo, RKO), and lung (H23, VMRC, H1299, A549) cancer cell lines, along with normal fibroblasts (F02, F03), as well as (E) *MYC*, mutant *KRAS*, and mutant *TP53* overexpressing K15 fibroblasts. Viability was assessed 48 h after transfection using the ATPlite assay. Statistical significance was determined by two-way ANOVA (uncorrected Fisher’s LSD) relative to the siRNA control. (F) Wound healing assay demonstrating changes in migratory capacity of PANC1, DLD1, H1299, and A549 cells after *CAMOR* depletion. Data were evaluated using one-way ANOVA (uncorrected Fisher’s LSD), *n* = 3–5. (G) Clonogenic assay showing the effect of *CAMOR* knockdown on colony-forming ability in PANC1, DLD1, H1299, and A549 cells. Statistical analysis was performed using one-way ANOVA (uncorrected Fisher’s LSD), n=3. (H) Chromatin immunoprecipitation-qPCR analysis of c-Myc and p53 binding to the *CAMOR* promoter in the indicated cell lines. The qPCR results were normalized to the level of IgG nonspecific antibody. (I) Diagram summarizing the proposed oncogenic role of *CAMOR*

*CAMOR* was found to be predominantly present in the nuclei of cancer cells (Suppl. Fig. 4C). Silencing of *CAMOR* using siRNA significantly reduced cell viability in 11 out of 12 analyzed cancer cell lines, having no significant effect on decreasing the viability of the normal fibroblasts (Fig. 4D). In K15 immortalized fibroblasts, knockdown led to a decrease in viability; however, a more pronounced reduction was observed when each of the oncogenes was overexpressed in (Fig. 4E), suggesting that the functional impact of *CAMOR* increases in the presence of the expressed oncogenes.

In further transformation-related phenotype tests, conducted in PANC1, DLD1, H1299, and A549 cell lines, *CAMOR* depletion impaired the migratory capacity (Fig. 4EF) and significantly reduced colony-forming ability in analyzed cell lines (Fig. 4G). Additionally, knockdown of *CAMOR* sensitized DLD1 and PANC1 cell lines to standard chemotherapeutical protocols used in these cancer types (Suppl. Fig. 4D). In A549 and H1299 cell lines, siRNA targeting *CAMOR* alone produced an excessive decrease in these cell lines viability, therefore the effect of adding chemotherapeutics agents could not be observed.

To determine which oncoprotein most strongly regulates *CAMOR* expression, we performed chromatin immunoprecipitation (ChIP) using antibodies against c-Myc (which is expected to directly bind to the promoter) and p53 (which in the mutant form more likely binds indirectly (41) (Fig. 4H). In DLD1 and PANC1 cells, which are driven by both oncogenes, c-Myc exhibited stronger promoter binding. Similarly, in H1299 cells, with absent *TP53* expression, only c-Myc binding was detected. In contrast, in the VMRC-LCD cell line, with mutant *TP53*, mutant p53 binding to the promoter was more prominent.

Given the robust effects of *CAMOR* depletion in cancer cells (Fig. 4I), we further assessed the impact of its overexpression (Suppl. Fig. 4E). In PANC1 and A549 cell lines, the overexpression led to increased migratory capacity (Suppl. Fig. 4F) and enhanced colony formation (Suppl. Fig. 4G). For *CAMOR*, GSEA revealed significant enrichment of gene sets associated with histone methyltransferase activity, N-acetyltransferase activity, and transcription regulator activity (Suppl. Fig. 4H). To further validate these findings, the expression of KMT2A, EIF4A1, and CHD1L1 was analyzed following *CAMOR* silencing. *CAMOR* depletion significantly reduced KMT2A transcript levels in both A549 and PANC1 cells, whereas the expression of EIF4A1 and CHD1L1 was not significantly affected (Suppl. Fig. 4I).

To investigate the mechanism of *CAMOR* action, we identified a partially overlapping protein-coding gene, *CARNMT1*, as a potential cis-target of this lncRNA (Fig. 5A). Data from our previous study (1) indicated, that *CARNMT1* expression decreased following knock-down of the oncogenic trio (Fig. 5B). Silencing of *CARNMT1* significantly impaired the viability of H23, A549, H1299, DLD1, HT29, PANC1, BxPC3, and K15 cells, while this effect was weaker than that of *CAMOR* silencing (Fig. 5C). Interestingly, the effects were not additive and a simultaneous silencing of both genes partially rescued the effects caused by the *CAMOR* depletion alone in A549, H1299, DLD1, HT29, BxPC3, and K15 cells. To further evaluate the roles of *CAMOR* and CARNMT1 in cell proliferation, we analyzed growth curves of A549 and PANC1 cells overexpressing either transcript. *CAMOR* significantly increased the proliferation rate in both cell lines, while CARNMT1 exerted a milder while significant positive effect only in PANC1 cells (Fig. 5D). These results suggested that effects of *CARNMT1* on cancer cell phenotype are weaker than of *CAMOR* and could interfere with pro-oncogenic activity of *CAMOR*.

**Figure 5.**
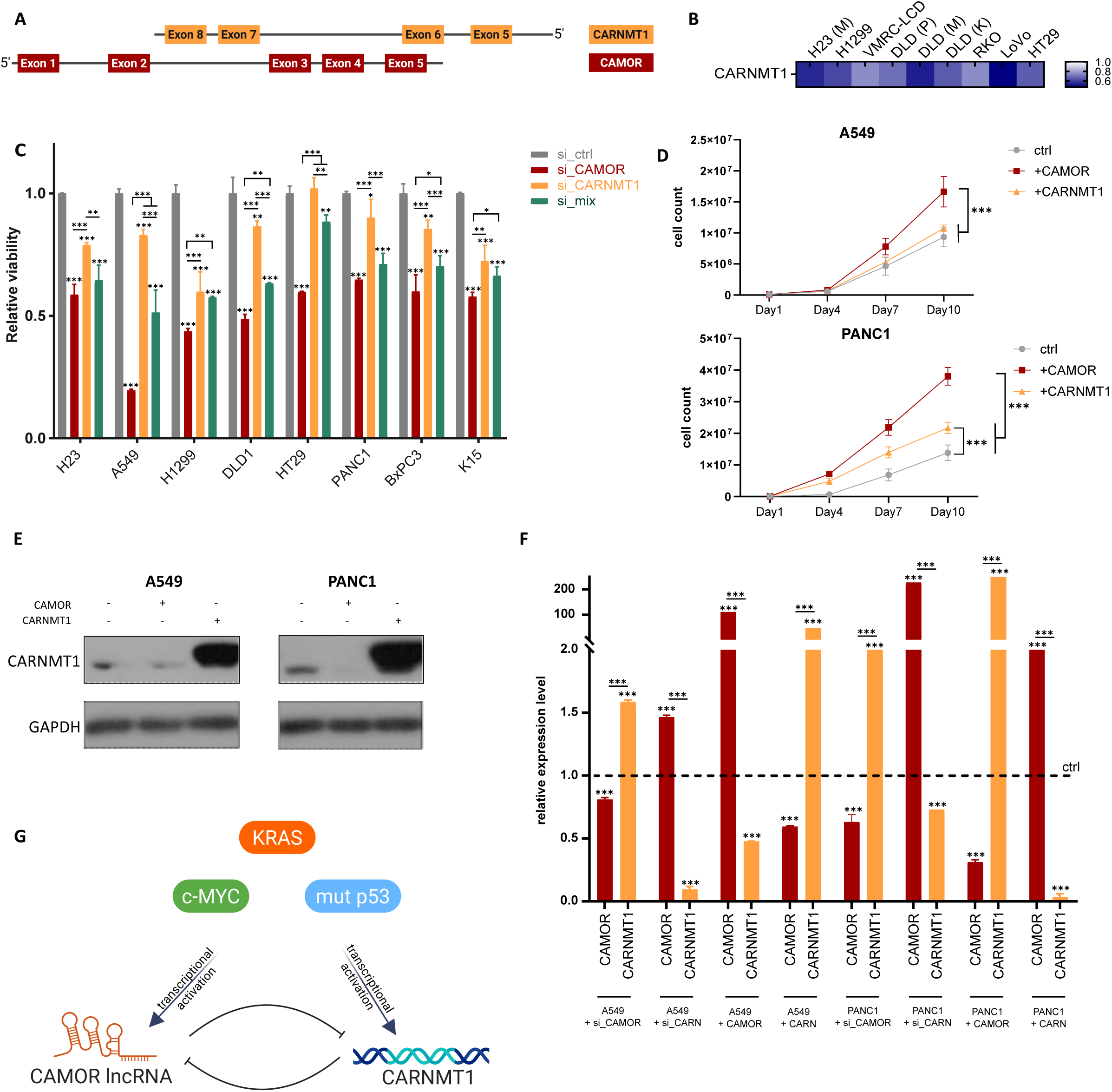
Regulatory feedback loop between *CAMOR* and CARNMT1. (A) Schematic depiction of the genomic localization and relative orientation of the *CAMOR* and *CARNMT1* genes. (B) Data from RNA-seq presenting changes in *CARNMT1* expression (fold change) following editing of *MYC*, mutant *KRAS*, and mutant *TP53.* (C) Viability of pancreatic (PANC1, BxPC3), colon (DLD1, HT29), and lung (H23, A549, H1299) cancer cell lines following silencing of *CAMOR* and/or *CARNMT1*. (D) Proliferation curves of A549 and PANC1 cells overexpressing *CAMOR* or *CARNMT1*. Data were evaluated using one-way ANOVA (uncorrected Fisher’s LSD), *n* = 3. (E) CARNMT1 protein levels in A549 and PANC1 cell lines upon *CAMOR* overexpression, as determined by western blot analysis. CARNMT1 and GAPDH were detected on separate membranes prepared in parallel from the same protein lysates. Equal amounts of protein were loaded onto each membrane. Full-length original blots are presented in Supplementary Figure 5. (F) Relative expression of *CAMOR* and *CARNMT1* following reciprocal silencing or overexpression in A549 and PANC1 cell lines. Data were evaluated using two-way ANOVA (uncorrected Fisher’s LSD), presented as means with SD, *n* = 3. (G) Proposed model illustrating a regulatory feedback loop between *CAMOR* and CARNMT1.

To assess the potential reciprocal regulation between CARNMT1 and *CAMOR*, we performed Western blot and expression analyses. Overexpression of *CAMOR* reduced CARNMT1 protein levels (Fig. 5E), suggesting negative regulation. Consistently, *CARNMT1* mRNA levels decreased upon *CAMOR* overexpression and increased following its silencing in A549 and PANC1 cells (Fig. 5F). Interestingly, *CARNMT1* overexpression led to a decrease in *CAMOR* expression, whereas *CARNMT1* silencing resulted in its upregulation. These findings support the existence of a negative feedback regulatory loop between CARNMT1 and *CAMOR* (Fig. 5G). However, a mild pro-proliferatory role of CARNMT1 is likely to cause a lack of its direct, clear anti-oncogenic effect via inhibition of *CAMOR* expression.

## DISCUSSION

To address current limitations in lncRNA research, which in most cases focuses on the clinical and biological significance of individual lncRNAs within a single cancer type (28, 42–46), we adopted an alternative approach. We integrated data from cancer cell lines derived from three distinct cancer types and analyzed them in the context of broad driver oncogene networks - *MYC* and oncogenic mutations of *KRAS* and *TP53*. Among these oncogenic drivers, *MYC* was associated with the highest number of lncRNAs exhibiting deregulated expression. Moreover, cluster analysis revealed that *MYC*–dependent lncRNA expression patterns were more strongly associated with this presence than with the tissue of origin of the respective cell lines. This is consistent with mostly non-redundant, broad transcriptional programs of c-Myc compared to mutant KRAS and mutant p53 programs found in different cell types (13).

We functionally validated two *MYC*-dependent lncRNAs, *AC104447.1* and *LINC00997*. Only *LINC00997* has been previously investigated in the context of cancer, although without the specific oncogene dependency. *LINC00997* has been reported to promote proliferation, migration, and metastasis in kidney renal clear cell carcinoma (47), cervical cancer (48), esophageal squamous cell carcinoma (ESCC) (49), and colorectal cancer (50). In our study, in colorectal cancer cells, migration was the only cellular parameter affected by *LINC00997* while in the pancreatic and lung cancer cell lines this lncRNA promoted cell viability, migration, and colony formation. Additionally, we found that serum and tumor tissue levels of *LINC00997* correlated in pancreatic cancer patients, confirming its potential use as a diagnostic biomarker suggested earlier for ESCC (49).

The second *MYC*–dependent lncRNA studied here - *AC104447.1*, as well as the analyzed mutant *KRAS*-dependent lncRNA *RAKRAR*, have not been previously described in the cancer context. Our functional validation indicates that these lncRNAs may possess broad pro-oncogenic potential, as they affected cell viability and colony formation in pancreatic, colorectal, and lung cancer cell lines, as well as cell migration in lung and colorectal (*AC104447.1*) and pancreatic (*RAKRAR*) cancer cell lines.

Only a few lncRNAs have a solid experimental evidence of oncogenic activity across three or more cancer types, with HOTAIR, MALAT1, MEG3, H19, GAS5, and NEAT1 being the most widely implicated (40). This, however, may reflect the fact that these lncRNAs were among the earliest discovered and most extensively studied. Nevertheless, our study strongly suggests existence of other such lncRNAs demonstrating pan-cancer functionality. Apart from the already mentioned examples the most consequent pan-cancer onco-lncRNA in our hands was *CAMOR*. Its silencing consistently reduced the viability of colon, lung, and pancreatic cancer cell lines, without affecting normal cells, suggesting a crucial role in tumor cell survival and positioning it as a potential therapeutic target in cancers driven by c-Myc, mutant KRAS, and mutant p53. In our recent study, we showed that even if a target gene is controlled by each of the three mentioned oncoproteins by redundancy, often a dominant may reduce an influence of the others due to competition (13). Consistently with this mechanism, we observed that c-Myc bound to the *CAMOR* promoter preferentially to mutant p53.

Anti-sense lncRNAs can modulate an expression of neighboring genes through *cis*-regulatory mechanism, typically affecting genes located in close genomic proximity (within 5kb) (51). In our study, we found that *CAMOR* lncRNA is localized in an anti-sense position to *CARNTM1*, and modulates its expression on the transcription level. CARNMT1 is a methyltransferase responsible for converting carnosine to anserine (52). Although its specific role in cancer remains unclear, CARNMT1 has pan-cancer presence, it has been identified as an HDAC1 substrate, and CARNMT1-dependent methylation of His residues present in C3H-type zinc finger domains modulated mRNA decay and alternative splicing (53, 54). Our observations indicate that CARNMT1 alone exerts a minor positive effect on cancer cell line viability, and it may mildly reduce but not significantly counteract the proliferative-promoting activity of *CAMOR*. This indicates a negative feedback regulatory interaction between the two transcripts, while the functional tests do not implicate a clear tumor suppressive activity of CARNMT1.

LncRNAs are known to regulate other genes at every stage of their expression. Since we observed that *CAMOR* affects both the transcript and protein levels of CARNMT1, and its subcellular localization is predominantly nuclear, its mechanism of action may involve epigenetic or transcriptional regulation. lncRNAs were described to recruit PRC2, DNMTs, or HDACs to a promoter or gene body, leading to increased H3K27me3 levels, DNA methylation, or histone deacetylation (55–57). At the transcriptional level, antisense lncRNAs can recruit repressive transcription factors (58), regulate splicing (59), or form R-loop, RNA-RNA duplex, or RNA-DNA triplex structures, thereby influencing transcription (60). Identification of the mechanisms by which *CAMOR* regulates CARNMT1 and vice-versa requires further investigation, while our study shows that it does not constitute a critical component affecting the oncogenic activity of *CAMOR*.

The GSEA provided insight into the potential molecular mechanisms underlying the oncogenic activities of the studied lncRNAs. Notably, the enriched gene sets differed between individual lncRNAs, suggesting that they may contribute to cancer cell phenotypes through distinct regulatory processes. For *AC104447.1*, the downregulation of ARHGAP5 expression following lncRNA depletion in pancreatic and lung cancer cells was consistent with its contribution to the enriched focal adhesion gene sets, supporting a potential regulatory role of *AC104447.1* in this pathway. Similarly, reduced CXCL13 expression following *RAKRAR* silencing in pancreatic and colon cancer cells, with a similar tendency in lung cancer cells, indicated the involvement of *RAKRAR* in adhesion-associated gene programs. For *CAMOR*, the significant downregulation of KMT2A following silencing in lung and pancreatic cancer cells was consistent with its association with epigenetic regulatory processes. In contrast, increased NDUFV2 expression following LINC00997 depletion did not support direct positive regulation of this gene, potentially reflecting a compensatory response to lncRNA loss.

Together, our findings provide experimental support for dependencies of the described lncRNAs on driver oncogenes which results in profound oncogenic phenotype alterations, reflected in pathways and genes regulated by lncRNAs. This highlights the multi-stage complexity and redundancy of the lncRNA-mediated regulation in cancer.

## STATEMENTS AND DECLARATIONS

## Author Contributions

Conceptualization, MG and DW; Methodology MG, DW, and AJ; Investigation, MG and AJ; Data curation, MG and AJ; Writing MG and DW; Funding Acquisition, MG and DW; Resources, MN-N, WK, TO; Supervision, MG and DW.

## Data Availability

The RNA-seq datasets analysed during the current study are available in the GEO database under accession number GSE239817. The original data are included in the article. For additional information, inquiries can be directed to the corresponding authors.

## Ethics approval

Fully anonymized patient sample collection and further laboratory experimental procedures were performed based on ethical committee approvals: No. 55/2023 of National Institute of Oncology in Warsaw and No. 109/2016 (with updates) of National Medical Institute of the Ministry of the Interior and Administration in Warsaw. Human primary fibroblasts (F02 and F03) were obtained from skin biopsies of healthy subjects based on bioethics committee approval (No. 108/2017 and 203/2020) of the National Medical Institute of the Ministry of the Interior and Administration in Warsaw. The study was conducted in accordance with the principles of the Declaration of Helsinki. Written consents for research use of the collected tissues were obtained from all patients.

## SUPPLEMENTARY FIGURE’S LEGENDS

**Supplementary Figure 1.** (A) Principal component analysis (PCA) of lncRNA expression from transcriptomics samples described in Fig. 1. Cell lines are indicated by colors and labels. (B) Transcript length-wise and (C) chromosomal-wise distribution of identified lncRNAs. (D) Average number of DE lncRNAs per downregulated oncogene. (E) Volcano plots of differentially expressed lncRNA groups for each downregulated oncogene. Upregulated lncRNAs are shown in red, and downregulated lncRNAs are shown in blue. Ribbon charts illustrating pools of DE lncRNAs dependent on *MYC*, mutant *TP53*, or mutant *KRAS* in colon (F) and lung (G) cancer cell lines. The left side shows the number of lncRNAs identified in cell lines with a single activated oncogene, whereas the right side presents the number of lncRNAs categorized as non-redundant (specific to a single oncogene), possibly redundant (shared with a co-expressed oncogene), or redundant (lncRNA pools overtaken by co-expressed oncogenes).

**Supplementary Figure 2.** qPCR validation of relative expression levels of (A) *LINC00997* and (B) *AC104447.1* following *MYC* silencing, compared to RNA-seq data. (C) Subcellular localization of *LINC00997* and (D) *AC104447.1* in the cytoplasmic and nuclear fractions of pancreatic (PANC1), colon (RKO), and lung (H1299) cancer cells. (E) Spearman correlation of *LINC00997* expression levels across pancreatic tumor tissues, normal tissues, and matched blood samples, measured by ddPCR (n=11). Functional enrichment analysis of upregulated genes associated with (F) *LINC00997* and (G) *AC104447.1* expression. The dot plot displays the top 10 enriched pathways/processes, ranked by the highest number of associated genes. The X-axis represents the Rich factor (% of Associated Genes), while the Y-axis indicates the enriched biological terms. The size of each dot corresponds to the number of genes mapped to the given pathway, and the color gradient represents the statistical significance based on the p-value, where red indicates higher statistical significance. Validation of expression of potential target genes following (H) *LINC00997* and (I) *AC104447.1* silencing. Data are presented as mean ± SD from *n* = 2 replicates per cell line and were analyzed using two-way ANOVA (uncorrected Fisher’s LSD) versus the siRNA negative control.

**Supplementary Figure 3.** (A) qPCR validation of *RAKRAR* expression following mutant *KRAS* silencing, compared to RNA-seq data; (B) Subcellular localization of *RAKRAR* in the cytoplasmic and nuclear fractions of pancreatic (PANC1), colon (DLD1), and lung (A549) cancer cells. (C) Functional enrichment analysis of upregulated genes associated with *RAKRAR* expression, shown as described in Suppl. Fig. 2F-G. (D) Validation of expression of potential target genes following *RAKRAR* silencing. Data are shown as mean ± SD from *n* = 2 replicates per cell line and were analyzed using two-way ANOVA (uncorrected Fisher’s LSD) versus the siRNA negative control.

**Supplementary Figure 4.** (A) qPCR-based confirmation of *CAMOR* expression changes following *MYC,* mutant KRAS, and mutant *TP53* silencing, consistent with RNA-seq results. (B) Spearman correlation of *CAMOR* expression levels across pancreatic tumor tissues, normal tissues, and matched blood samples, measured by ddPCR (n=9). (C) Distribution of *CAMOR* between nuclear and cytoplasmic compartments in pancreatic (PANC1), colon (DLD1), and lung (H1299, A549) cancer cells. (D) Cell viability of DLD1 and PANC1 cell lines following CAMOR knockdown in combination with standard chemotherapeutic treatments. (E) qPCR confirmation of *CAMOR* overexpression in A549 and PANC1 cell lines; (F) Migration and (G) colony formation assays assessing the functional effects of *CAMOR* overexpression in PANC1 and A549 cells. (H) GSEA of upregulated genes associated with *CAMOR* expression, highlighting enriched functional categories, shown as described in Suppl. Fig. 2F-G. (I) Expression analysis of selected genes (KMT2A, EIF4A1, and CHD1L1) following *CAMOR* silencing. Data are presented as mean ± SD from n = 2 replicates per cell line and were analyzed by two-way ANOVA with uncorrected Fisher’s LSD test versus the siRNA negative control.

**Supplementary Figure 5.** Uncropped blots for CARNTM1 and GAPDH

## SUPPLEMENTARY TABLE’S LEGENDS

**Supplementary Table 1.** RNA sequencing DElncRNAs analysis

**Supplementary Table 2.** Fold changes and validation of the lncRNAs with the lowest average upon CRISPR-Cas9-mediated disruption of (1) *MYC*, (2) mutant *KRAS*, and (3) *MYC*, mutant *KRAS*, and mutant *TP53*.

**Supplementary Table 3.** List of (1) primers sequences, (2) siRNAs sequences, and (3) antibodies used in this study.

