## Supplementary figures and images for "*CAMOR* and specific oncogene-driven lncRNAs mediate carcinogenic functions downstream of *MYC*, mutant *KRAS*, and mutant *TP53*"

### Suppl Figures

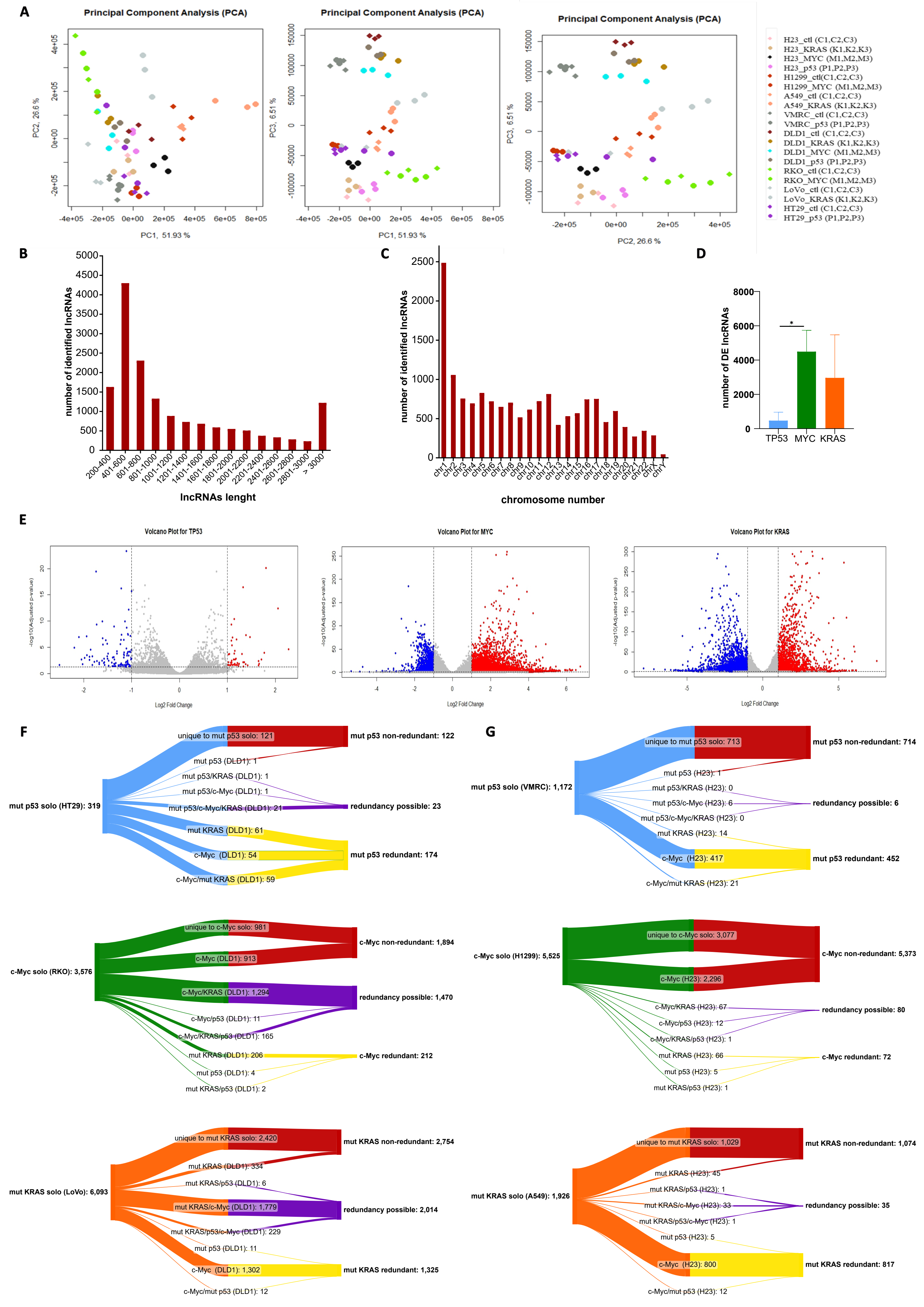

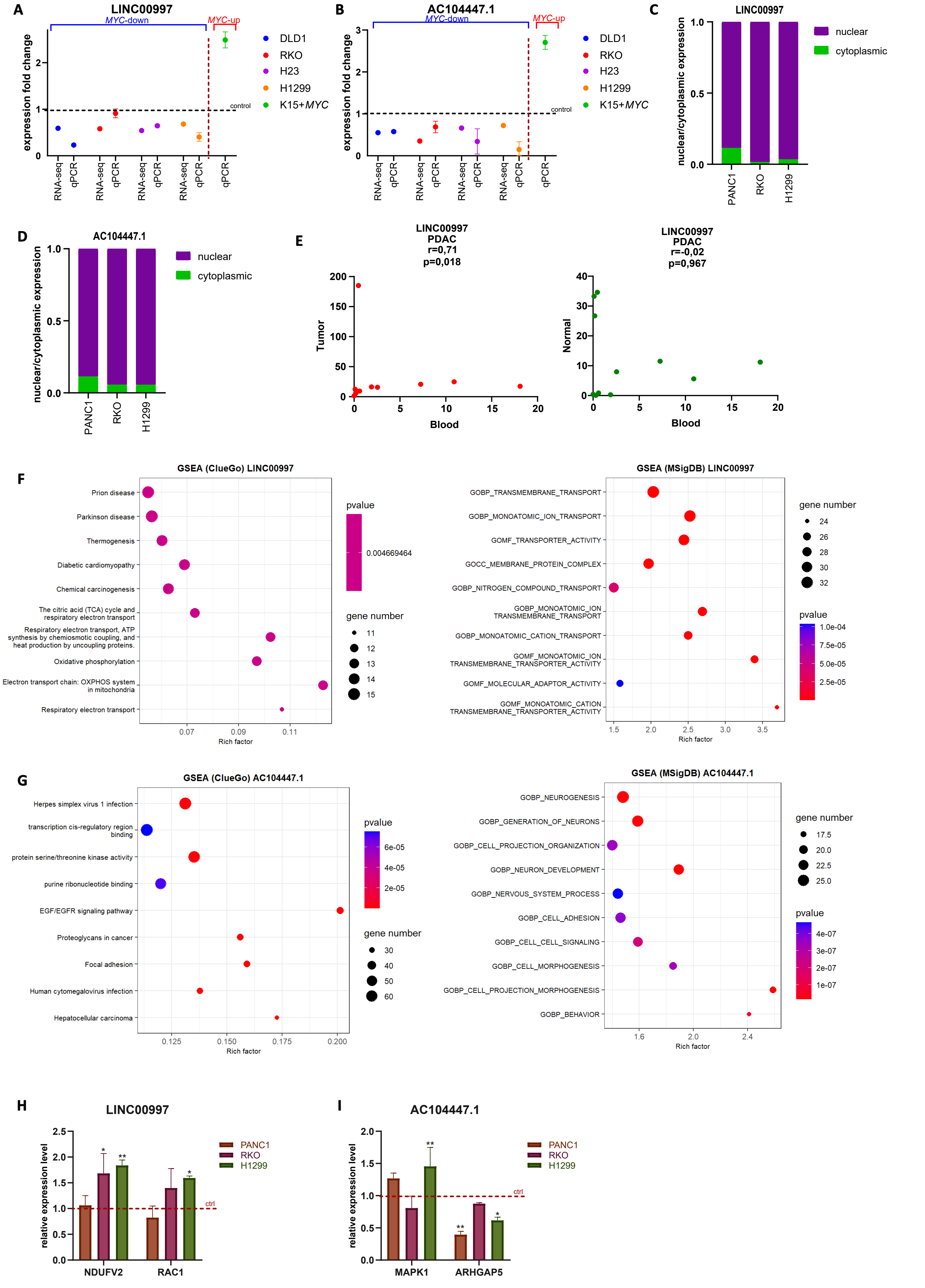

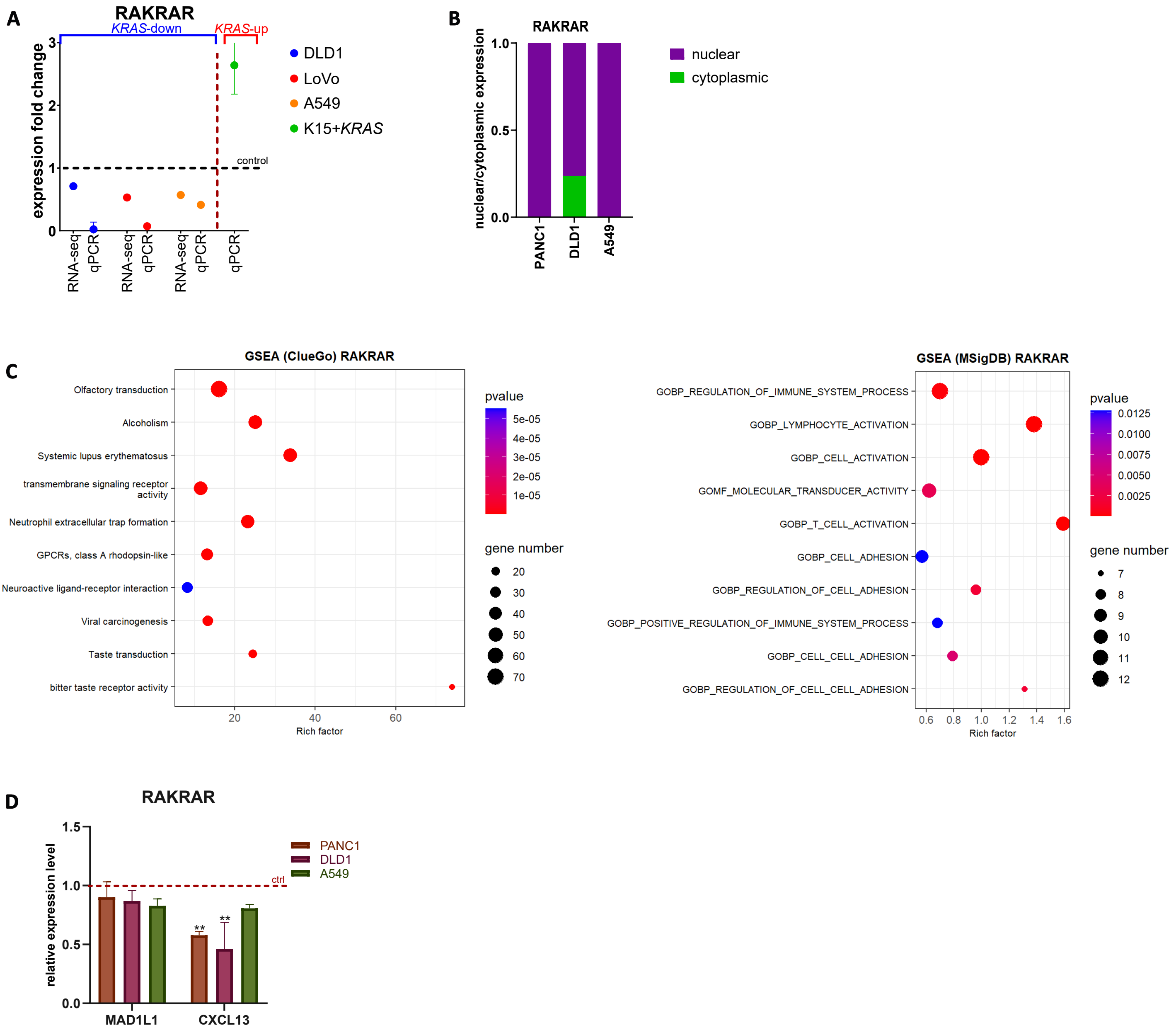

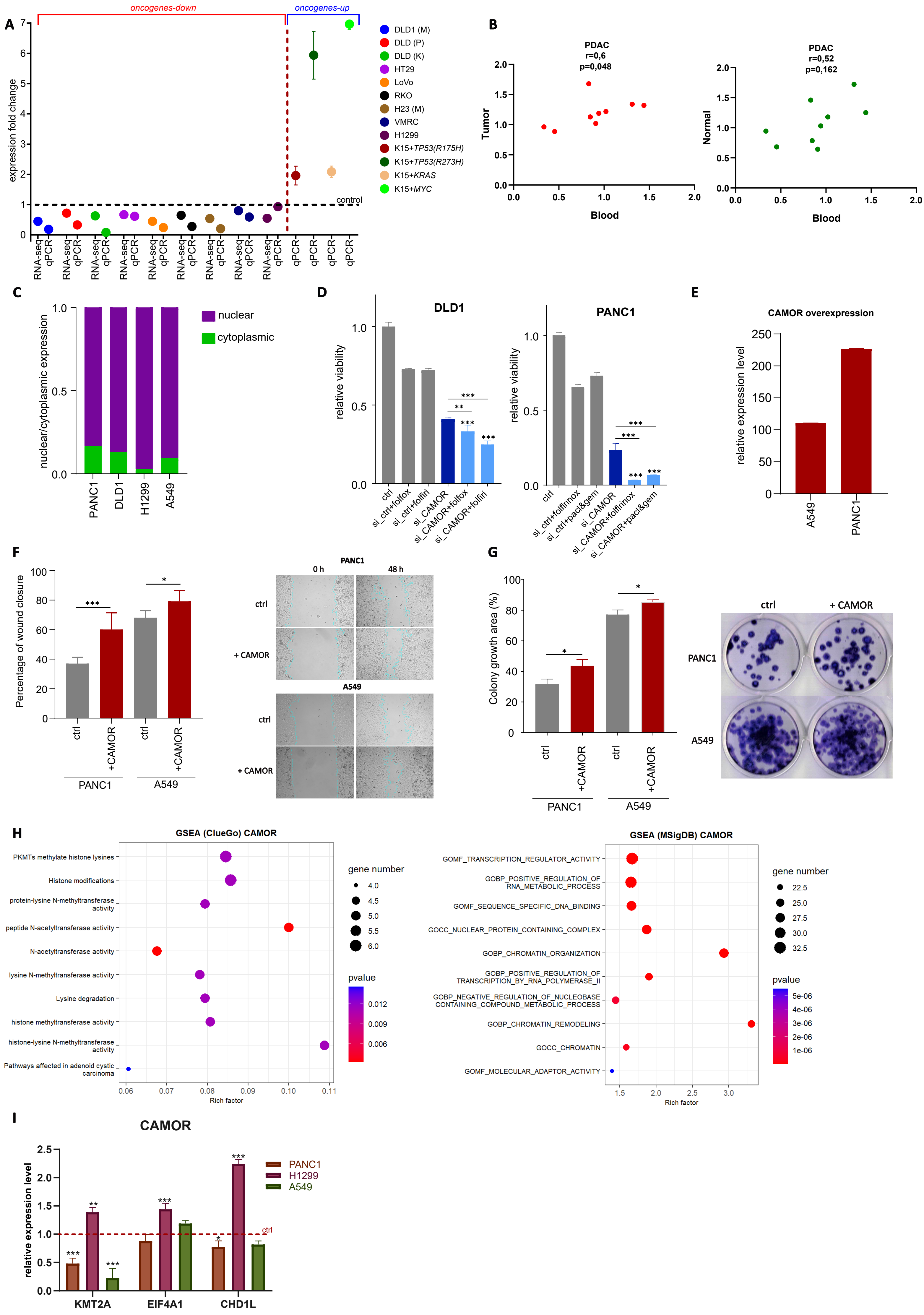

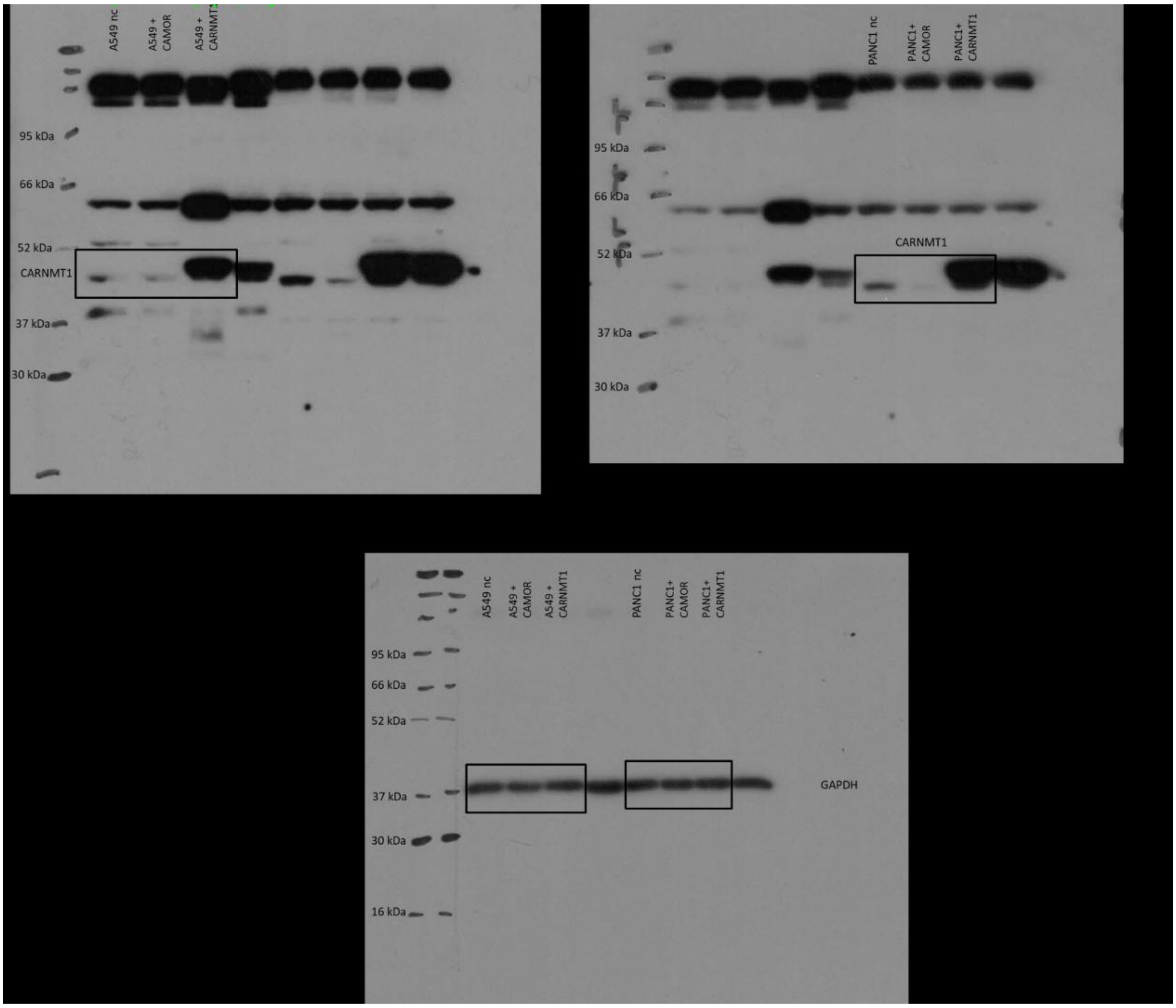
